# Photon-counting cine-cardiac CT in the mouse

**DOI:** 10.1101/660100

**Authors:** Darin P. Clark, Matthew Holbrook, Chang-Lung Lee, Cristian T. Badea

## Abstract

The maturation of photon-counting detector (PCD) technology promises to enhance routine CT imaging applications with high-fidelity spectral information. In this paper, we demonstrate the power of this synergy and our complementary reconstruction techniques, performing 4D, cardiac PCD-CT data acquisition and reconstruction in a mouse model of atherosclerosis, including calcified plaque. Specifically, *in vivo* cardiac micro-CT scans were performed in four ApoE knockout mice, following their development of calcified plaques. The scans were performed with a prototype PCD (DECTRIS, Ltd.) with 4 energy thresholds. Projection sampling was performed with 10 ms temporal resolution, allowing the reconstruction of 10 cardiac phases at each of 4 energies (40, 3D volumes per mouse scan). Reconstruction was performed iteratively using the split Bregman method with constraints on spectral rank and spatio-temporal gradient sparsity. The reconstructed images represent the first *in vivo*, 4D PCD-CT data in a mouse model of atherosclerosis. Robust regularization during iterative reconstruction yields high-fidelity results: an 8-fold reduction in noise standard deviation for the highest energy threshold (relative to algebraic reconstruction), while absolute spectral bias measurements remain below 13 Hounsfield units across all energy thresholds and scans. Qualitatively, image domain material decomposition results show clear separation of iodinated contrast and soft tissue from calcified plaque in the *in vivo* data. Quantitatively, spatial, spectral, and temporal fidelity are verified through a water phantom scan and a realistic MOBY phantom simulation experiment: spatial resolution is robustly preserved by iterative reconstruction (10% MTF: 2.8-3.0 lp/mm), left-ventricle, cardiac functional metrics can be measured from iodine map segmentations with ∼1% error, and small calcifications (615 μm) can be detected during slow moving phases of the cardiac cycle. Given these preliminary results, we believe that PCD technology will enhance dynamic CT imaging applications with high-fidelity spectral and material information.

## Introduction

Dual-energy (DE, spectral) imaging methods enhance the diagnostic capabilities of x-ray CT through quantitative material discrimination [1] and the ability to synthesize virtual nonenhanced [2, 3] and monochromatic images [4]. Clinically, DE-CT scanners are available from several vendors: Siemens (dual-source), GE (kVp switching), and Philips (dual-layer). These DE scanners are associated with several routine imaging applications, including plaque differentiation, myocardial perfusion, and kidney stone characterization [5]. Preclinically, spectral CT has been utilized in several additional applications, including differential imaging of vasculature and vascular permeability in sarcoma [6] and lung [7] tumors, and with several preclinical contrast agents based on iodine [8], barium [9], gadolinium [10], and gold [6].

Unfortunately, the potential of spectral CT is limited by the energy-integrating detectors (EIDs) used by current CT systems. EIDs record a signal proportional to the detected photon flux, weighted by the photon energy and integrated across the entire energy spectrum. Thus, EID spectral resolving power is limited when used with polychromatic x-ray sources. Alternatively, photon-counting detectors (PCDs), which are currently under active development, promise to enable more robust spectral separation in multi-energy CT imaging applications [11, 12]. Ideally, photon counting improves spectral separation by binning incoming photons into two or more energy bins, improving material discrimination with a single detector and CT scan. The other potential advantages of PCD-CT include reduced electronic noise, higher contrast-to-noise ratios, improved spatial resolution, and improved dose efficiencies with enhanced clinical applications such as breast, temporal bone, and lung imaging [13].

Because PCD technology provides spectral separation with a single detector, its maturation promises to enhance routine CT imaging applications with high-fidelity spectral information. This enhancement has already been demonstrated in dynamic large animal [14] and rabbit [15] studies using prototype clinical hardware. Here, we demonstrate the potential of this synergy in small animal micro-CT, performing 4D (3D+time) PCD-CT data acquisition and reconstruction in a mouse model of atherosclerosis, without a significant increase in imaging dose relative to our 4D EID-CT imaging protocols.

To perform robust, 4D PCD-CT reconstruction, we refine and extend our multi-channel reconstruction framework based on the split Bregman method with the add-residual-back strategy [16] and on a low rank and sparse signal model [17]. We initially proposed this framework for 4D, DE CT reconstruction using EIDs [18]. We later extended this framework for spectral CT reconstruction with an arbitrary number of spectral channels, proposing a regularizer called rank-sparse kernel regression (RSKR) [9]. In this paper, we combine our original 4D, DE reconstruction framework with RSKR, enabling *in vivo*, 4D PCD-CT reconstruction. We show that the high fidelity of our reconstruction results enables simultaneous material characterization and dynamic imaging, without a significant increase in imaging dose relative to EID-based, preclinical cardiac micro-CT.

## Materials and methods

### System description and image acquisition

Our PCD micro-CT system uses a SANTIS 0804 ME prototype PCD (DECTRIS Ltd., Baden-Dättwil, Switzerland; www.dectris.com). The detector is constructed with a 1 mm thick CdTe sensor, 150-μm isotropic pixels (515×257), and four independent and fully adjustable energy thresholds (here, set to 25, 34, 40, 55 keV). The data were acquired with a G297 x-ray tube (Varian Medical Systems, Palo Alto, CA; fs = 0.3/0.8 mm; tungsten rotating anode; filtration: 0.1 mm Cu; 80 kVp, 5mA, 10 ms exposure/projection). The source-to-detector and source-to-object distances were 831 mm and 680 mm, respectively. To minimize ring artifacts in our reconstructions, we scanned using a helical trajectory with 3 rotations and 1.25 cm of total translation during scanning (30 seconds/rotation), acquiring a total of 9000 projections per threshold. The absorbed radiation dose associated with imaging was ∼190 mGy (vs. ∼170 mGy for our 4D EID-CT imaging protocols [19]). These doses are comparable to many commercial micro-CT scanners when reconstructing 10 cardiac phases [20], and are consistent with our previous results showing dose benefits associated with four-threshold, PCD-based spectral imaging over DE EID-CT [21]. Notably, additional PCD dose optimizations (e.g. threshold and kVp selection, beam filtration, etc.) are likely possible for 4D imaging and are still under investigation.

The animal scanning in this work was performed following a protocol approved by the Duke University Institutional Animal Care and Use Committee. The four mice scanned were female ApoE-/-mice. ApoE-/-mice have germline deletion of the apolipoprotein E gene [22], show a marked increase in total plasma cholesterol levels, and are prone to develop atherosclerotic lesions [23]. At 8-12 weeks of age, the ApoE-/-mice were exposed to 25 fractions of 2 Gy partial-heart irradiation, mimicking cardiac radiation exposure of breast cancer patients treated with radiation therapy [24]. After irradiation, the mice were kept on a regular diet for a year prior to imaging. Three days before imaging, the mice were intravenously injected with gold nanoparticle contrast agent (15 nm *AuroVist*, www.nanoprobes.com) at a dose of 0.004 mL/g mouse, with gold accumulation expected at the site of any myocardial injury [24]. Notably, with an 80 kVp source spectrum, K-edge imaging of gold (edge at 80.7 keV) was not possible. Three days later, immediately prior to imaging, the same mice were injected with a liposomal iodinated contrast agent (described in [8]; 0.012 mL/g mouse) to measure cardiac functional metrics. The free-breathing mice were scanned while under anesthesia induced with 1-2% isoflurane delivered by nose cone. Both ECG and respiratory signals were recorded during scanning (average heart rate: 448 ± 50 bpm; average respiratory rate: 142 ± 19 breaths/min.). The ECG signal was used for retrospective projection weighting during reconstruction. Respiratory gating was not performed.

### Multi-channel image reconstruction

Following from our past work on data-adaptive, iterative reconstruction methods for temporal and spectral CT [9, 18, 25], Fig 1 summarizes our approach to these multi-channel reconstruction problems. We solve a series of weighted algebraic reconstruction problems (columns of X, channels indexed by *c*), subject to relaxed constraints which ensure consistency between channels (“Reg”; scalar regularization parameter, λ), using the split Bregman method with the add-residual-back strategy [16] and a low-rank and sparse signal model [17]. During initialization, an initial estimate is produced for each channel (step 1). In this work, the weights for each channel, *Q*_*c*_, select for energy (y, vectorized, log-transformed projection data for all thresholds) and assign weights to projections based on the cardiac phase during which they were acquired relative to the phase being reconstructed [18]. The regularization parameter for each channel, *μ*_*c*_, is scaled based on the data and a vector of scalar multipliers, *α*, which can account for differential noise levels between input channels (step 2). Iterative reconstruction (steps 3-5) then proceeds with consecutive regularization (step 3), residual update (step 4), and data fidelity update (step 5) steps, as in the split Bregman method [16].

**Fig 1.**
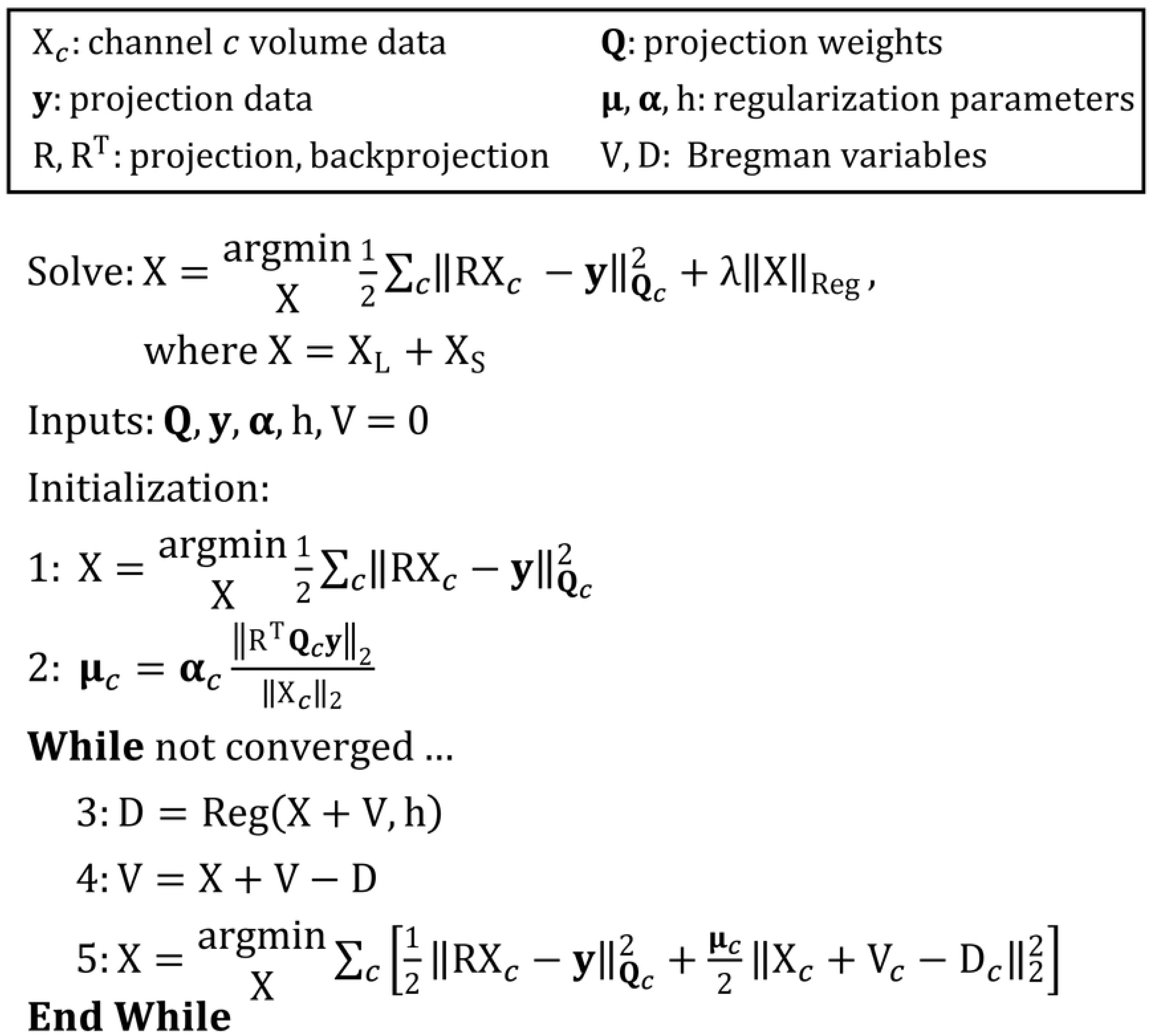
Generalized, multi-channel CT reconstruction framework. Indexing channels by *c*, the method solves for a series of reconstructed volumes, X_*c*_, subject to one (or more) multi-channel regularizers (“Reg”).

While many forms of regularization are possible within this robust framework, here, we employ two multi-channel regularizers (as detailed in previous work [26]): (1) RSKR along the energy dimension to enforce low spectral rank across all cardiac phases simultaneously, and (2) 4D, joint bilateral filtration to enforce consistent spatio-temporal gradient sparsity patterns between energies. Relative to Fig 1, the split Bregman method readily extends to handle these two multi-channel regularizers, with independent V (residual) and D (regularization) terms for each regularizer and an additional L2 term in step 5.

Notably, the projections for each PCD energy threshold were not subtracted to compute energy bin projections, as subtraction amplified noise prior to regularized reconstruction and was not required for material decomposition. In total, we simultaneously reconstructed 40 volumes (360×360×224 voxels/volume; 10 cardiac phases × 4 energy thresholds) with 123-μm, isotropic voxels. Reconstruction was performed using our custom GPU-based, multi-channel reconstruction toolkit [27] and four NVIDIA Titan Xp GPUs on an Ubuntu Linux workstation with 256 GB of system RAM and two Intel Xeon E5-2650 processors. The total reconstruction time was ∼400 minutes (∼10 minutes/volume; initialization + three Bregman iterations).

### Material decomposition

Extending the approach of Alvarez and Macovski [28], we performed image-domain material decomposition:

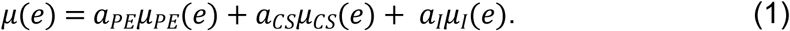

Given reconstructions for each energy threshold, the first two terms describe the energy-dependent attenuation, *µ*(*e*), owing to the photoelectric effect (PE) and Compton scattering (CS), each normalized to one in water. The last term is a basis function for iodine (I), whose K-edge is bracketed by the chosen detector thresholds (25, 34, and 40 keV). Gold, which accumulated in the liver and spleen, appears mainly in the PE component because the 80 kVp source spectrum did not include the K-edge of gold (80.7 keV).

Following previous work [6, 21], material decomposition was calibrated by scanning a 3D-printed, physical calibration phantom using data acquisition parameters identical to the *in vivo* data. The phantom has similar dimensions to a mouse and cradle (3.8 cm diameter) and is made from the same material as the mouse cradle used during CT scanning (polylactic acid, PLA). The phantom contained vials of water and reference concentrations of iodine and calcium in water. Linear regression was used to determine the sensitivity of iodine, and a singular value decomposition was used to more closely fit the PE and CS basis functions to the measured attenuation values for PLA, calcium, and water. These sensitivity measurements were arranged in a sensitivity matrix (*µ*_*PE*_, *µ*_*CS*_, *µ*_*I*_ per energy threshold in Eq 1) to perform material decomposition (solve for *a*_*PE*_, *a*_*CS*_, *a*_*I*_ per voxel) by matrix pseudo-inversion subject to a non-negativity constraint.

For reference, following unit-vector normalization per material, the condition number of the calibrated material sensitivity matrix was 45. Higher condition number values indicate greater potential for error amplification. This decomposition was more poorly conditioned than our previous two-material, EID-based decompositions, e.g. the separation of gold and iodine (5.6) [6], but better conditioned than our previous decomposition of four-threshold PCD data into PE, CS, iodine, and barium maps (∼100) [9].

### Quantitative analysis

To better understand the limitations of our reconstruction method, we first performed a simulation experiment using the MOBY mouse phantom [29]. Following past work [18], the phantom was constructed at 100 subphases to model temporal blurring in reconstructing 10 phases of the cardiac cycle (respiratory motion was not modeled). Two, 5×5×5 voxel (615-μm length) calcifications were attached to the left ventricle to assess motion and size artifacts resulting from data acquisition and reconstruction. The phantom’s material composition was parameterized using the experimentally calibrated sensitivity matrix, and projections (quantized to the nearest subphase) were digitally acquired using the ECG signal from the first mouse (the mouse with the highest heart rate, 489 bpm) and our GPU-based reconstruction toolkit [27]. Poisson noise was added to the projection data to approximately match *in vivo* noise levels; however, PCD-based spectral distortions were not modeled.

To demonstrate the fitness of the simulated and *in vivo* PCD data for calcified plaque and cardiac functional analysis, we segmented the left ventricle of the heart and the calcified plaque using the material decomposition results and the Avizo (v9.2) and ITK-SNAP [30] (www.itksnap.org) software packages. Left ventricle volume curves were used to determine end-systolic volume (ESV) and end-diastolic volume (EDV) measurements, which were then used to calculate stroke volume (SV = EDV − ESV), ejection fraction (EF = SV/EDV), and cardiac output (CO = SV * heart rate).

To supplement the *in vivo* results, we further characterized our PCD-based micro-CT system and reconstruction algorithm with modulation transfer function (MTF) and noise power spectrum (NPS) measurements. Specifically, we acquired PCD-based micro-CT data of a water cylinder (3 cm diameter) using data acquisition parameters identical to the *in vivo* data. We reconstructed the water cylinder exactly as the *in vivo* data, using the ECG signal from the first mouse for gating. 64×64×64 voxel volumes of interest (VoIs) around the periphery of the water phantom and at three different z positions were then used to measure the NPS for each PCD threshold setting (Eq 4.1 in this reference [31]; aggregated over all 10 phases; 960 total VoIs averaged / threshold). Radial line profiles from the center of the phantom were used to derive the edge spread function and then the axial MTF for each PCD threshold [31] (derived from 32 line profiles measured on the temporal-average volume).

## Results

### Simulation experiment

Fig 2 summarizes the results of the PCD-CT, MOBY phantom simulation experiment (results shown in a 2D region of interest around the heart). Our multi-channel reconstruction approach significantly reduces the noise at the highest energy thresholds (40-80 keV; 55-80 keV), while preserving details from the lowest energy threshold (25-80 keV; A vs. B). Because the ground truth reconstruction is known in this simulation experiment (C), root-mean-square error (RMSE; Hounsfield units, HU) measurements can be computed over the entire reconstruction (global, white text) and over only the portion of the reconstruction which changes in time (temporal, yellow text). Our data-adaptive iterative reconstruction (B) roughly equalizes the reconstruction error across all time points and energies, despite a nearly 5-fold difference in reconstruction errors between the lowest and highest energy threshold following algebraic reconstruction (A). The accuracy of these reconstruction results is consistent with our previous, detailed analysis of spectral bias introduced by RSKR and our iterative reconstruction approach [9].

**Fig 2.**
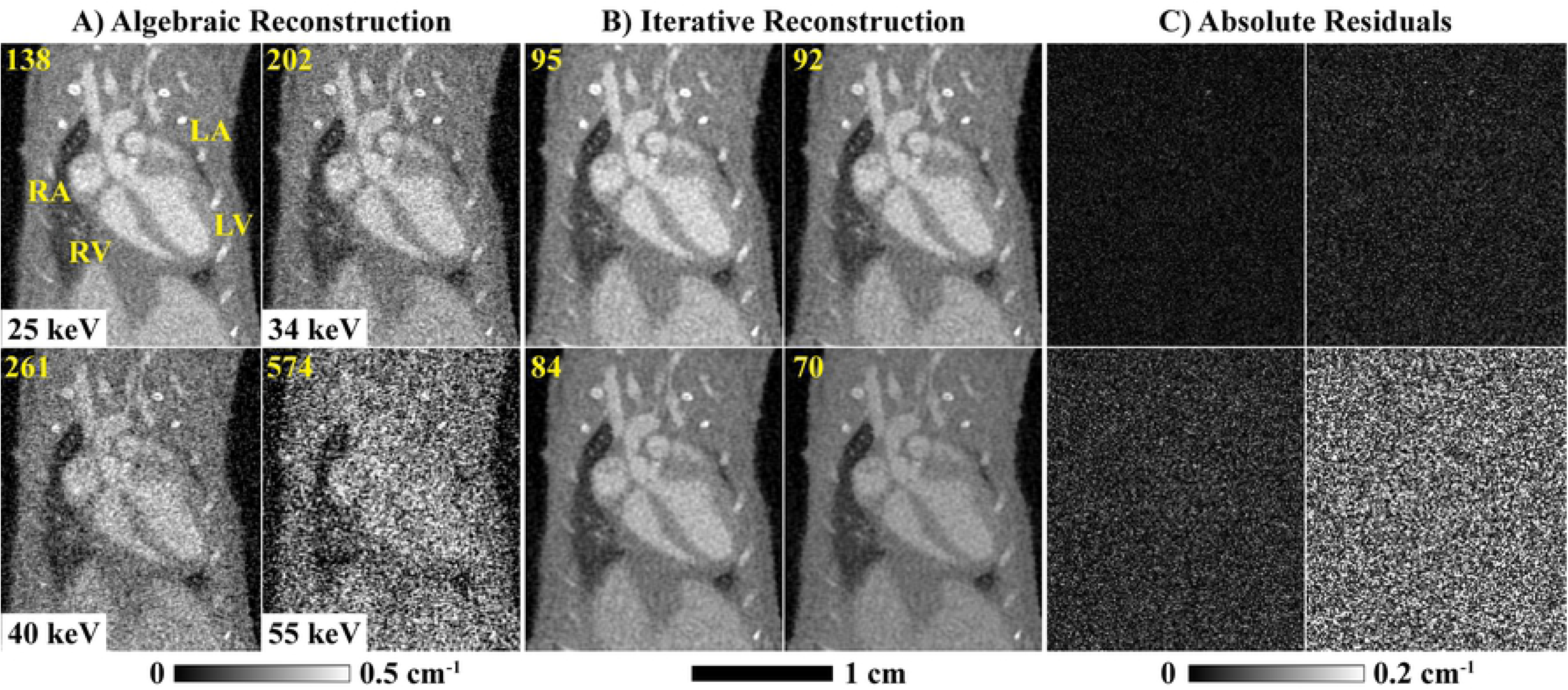
3D+time+energy MOBY phantom simulation results. Using *in vivo* ECG gating data to create projections, reconstruction results are shown at ventricular diastole following algebraic reconstruction (A, four thresholds; Fig 1, step 1) and iterative reconstruction (B; Fig 1, steps 3-5) vs. the expected reconstruction results (C). Global (white) and temporal (yellow) RMSE results (HU) are shown at the top of each result and are computed over all cardiac phases. The results of material decomposition (D, iterative reconstruction, four cardiac phases) are compared with the expected results (E) and the absolute residuals (F). Material color-coded text denotes the global (left) and temporal (right) RMSE values for each material map (D) as well as the phantom’s material composition (E). White and blue arrows highlight features referred to in the text. Cardiac labels (yellow text): LV, left ventricle; RV, right ventricle; LA, left atrium; RA, right atrium.

Fig 2 D-F compare the obtained and expected material decomposition results, with global (left) and temporal (right) RMSE values indicated for each material. As mentioned, the phantom contains two, 5×5×5 voxel calcifications. The top calcification, which moves 6 times its width over the cardiac cycle, is seen to disappear in the material decomposition results obtained by iterative reconstruction (white arrows). The bottom calcification, which moves 2.5-times its width over the cardiac cycle, is resolved in all 4 cardiac phases shown; however, partial volume effects reduce the apparent size of both calcifications (blue arrows).

Despite limitations associated with fast-moving calcifications, Fig 3 affirms that our iterative reconstruction and material decomposition methods yield robust left-ventricle (LV) cardiac functional metrics. The percent error (100% * |expected − measured|/ expected) in the LV volume measurements is less than 6% at every phase of the cardiac cycle. Small volume measurement errors at end diastole and end systole lead to ∼1% error in the derived cardiac functional metrics, even though this simulation experiment was conducted using the ECG signal from the mouse with the fastest heart rate (489 beats/min.). Combined with our previous experimental results for dual-energy, cardiac CT [18], these simulation results suggest that our iterative reconstruction method yields robust *in vivo* LV volume measurements and derived functional metrics.

**Fig 3.**
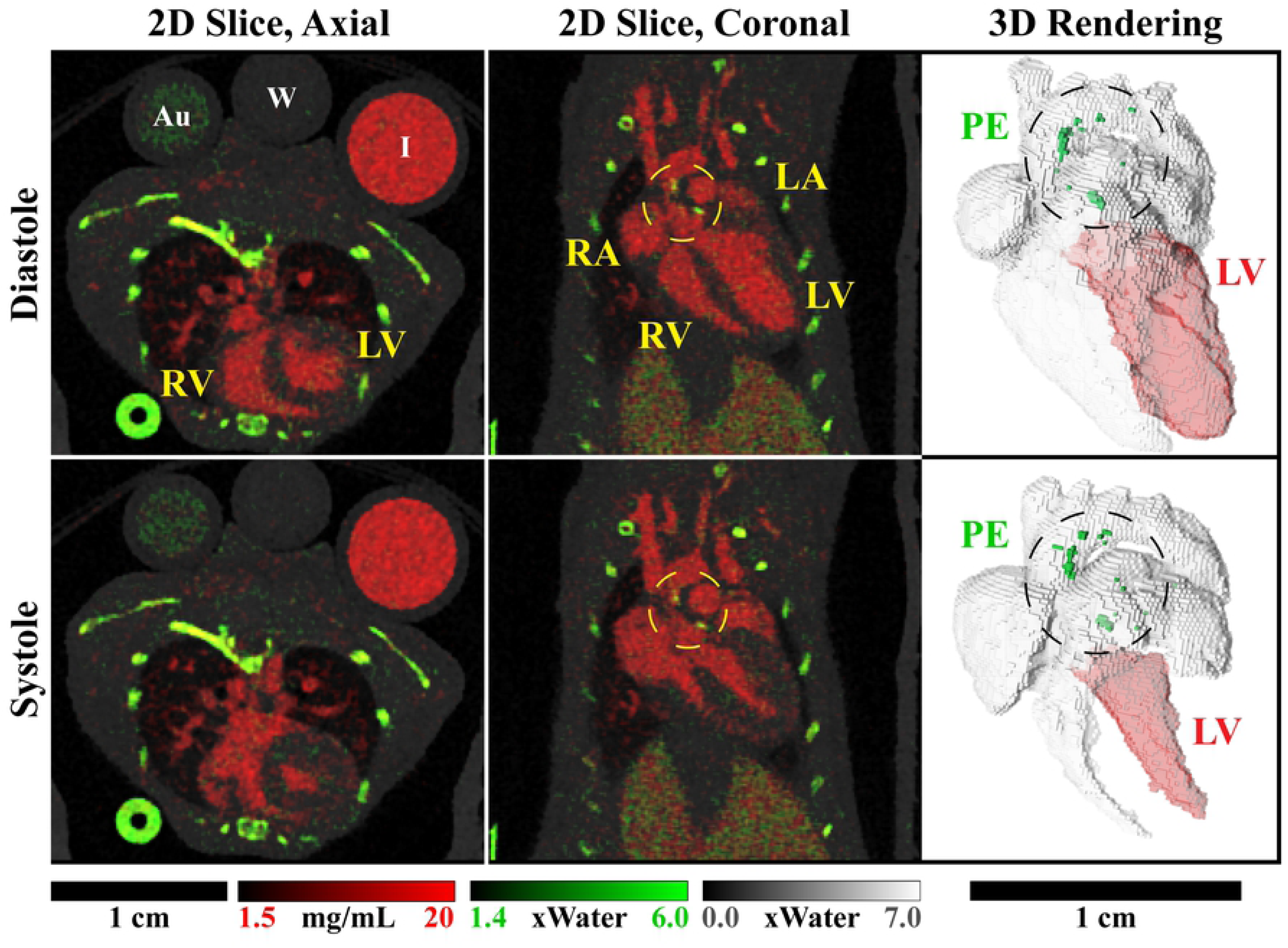
Cardiac functional measurement accuracy in the MOBY phantom simulation experiment. Expected LV volume curves measured in the reference phantom (black line; Fig 2E) are compared with experimentally measured curves segmented using the iodine map following iterative reconstruction (red line; Fig 2D). Derived LV functional metrics are compared within the plot’s legend: HR, simulated heart beats/minute; SV, stroke volume (μL); EF, ejection fraction; CO, cardiac output (mL/minute).

### System characterization

Fig 4 summarizes the MTF (A) and NPS (B) measurements taken in the physical water phantom, following data acquisition and reconstruction consistent with the *in vivo* data. The MTF curves are seen to shift to higher spatial frequencies between the initial algebraic reconstruction results (“Init.”) and the final iterative reconstruction results (“Iter.”), with the 10% MTF intersection increasing from 2.2-2.4 line pairs / mm (lp/mm; depending on threshold, “T”) to 2.8-3.0 lp/mm. This increase is not surprising given additional data fidelity updates during iterative reconstruction, which better resolves high spatial frequencies, and given the piece-wise constant signal model associated with regularization based on bilateral filtration, which tends to quantize partial volume effects. Consistent with neighborhood-based regularizers, the noise power is seen to decrease significantly at high spatial frequencies and more moderately at low spatial frequencies, following iterative reconstruction. Notably, however, the data-adaptive nature of the reconstruction algorithm very successfully matches noise properties between energies.

**Fig 4.**
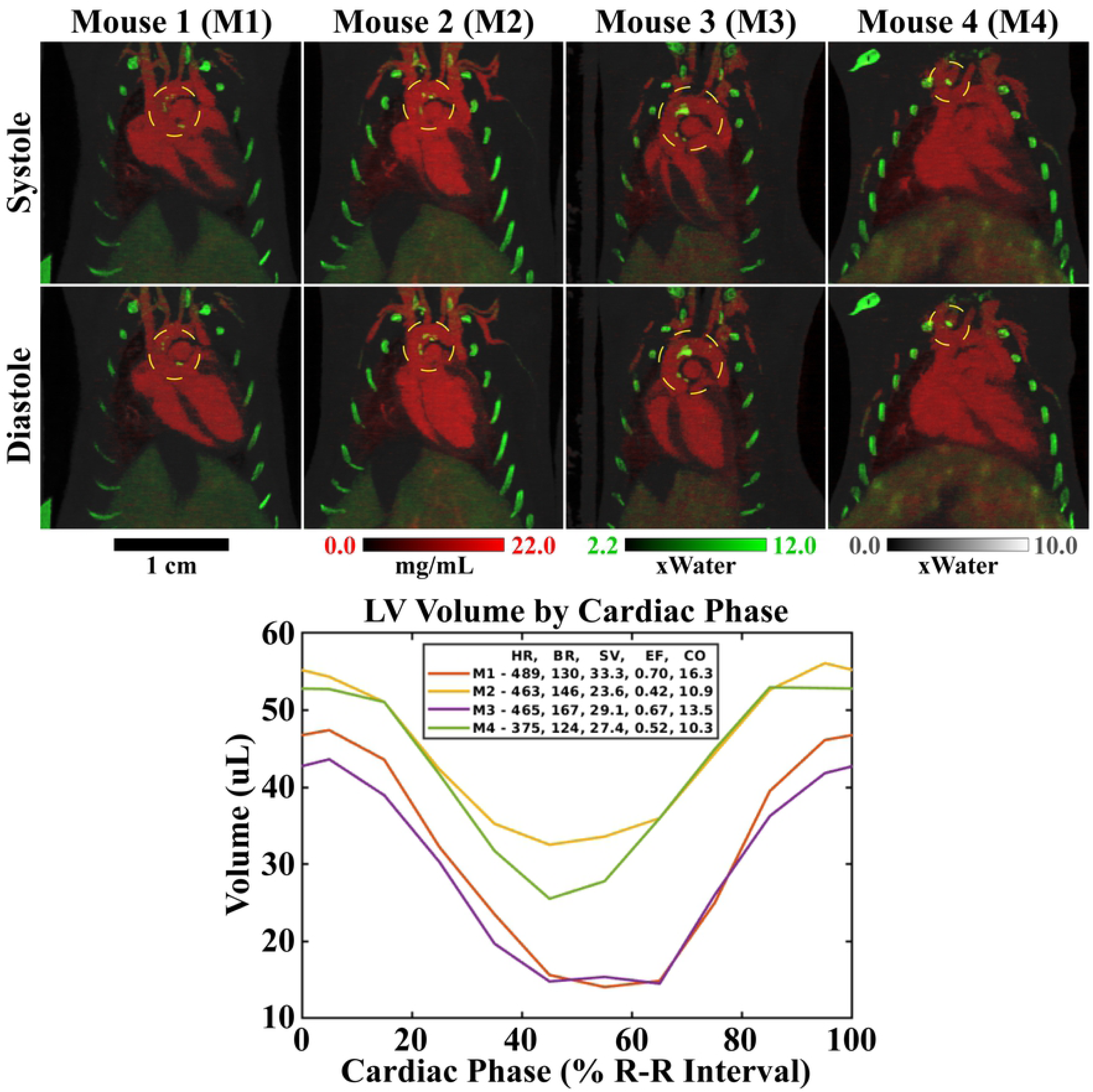
Axial MTF (A) and NPS (B) measurements. Measurements were taken in a 3 cm, physical water phantom after algebraic reconstruction (“Init.”) and after iterative reconstruction (“Iter.”). Cardiac gating of the phantom projection data was artificially performed using the ECG signal from the mouse with the highest heart rate (489 beats/min.). Results are plotted per energy threshold (T1, 25-80 keV; T2, 34-80 keV; T3, 40-80 keV; T4, 55-80 keV). 10% MTF measurements are shown in the legend, in parentheses and in lp/mm.

### *In vivo* experiments

Fig 5 summarizes the results of multi-channel, iterative reconstruction applied to the *in vivo*, 4D, PCD-CT data. Specifically, the algebraic initialization (A) and iterative reconstruction results (B) are shown for each energy threshold setting (25-80 keV, 34-80 keV, 40-80 keV, and 55-80 keV) at ventricular diastole. Yellow labels indicate the four chambers of the heart in coronal orientation: left ventricle (LV), right ventricle (RV), left atrium (LA), and right atrium (RA). Consistent with the noise standard deviation values reported in yellow text (measured in the water vial; see Fig 6), the absolute residual images shown in (C), and the previous simulation results (Fig 2), our data-adaptive regularization scheme preserves the spatial and temporal resolution of the highest-fidelity threshold setting (25-80 keV) without compromising the spectral contrast seen with the more photon-starved threshold settings (40-80 keV and 55-80 keV). The effectiveness of our method is starkly illustrated by an 8-fold reduction in the noise standard deviation in water for the highest energy threshold.

**Fig 5.**
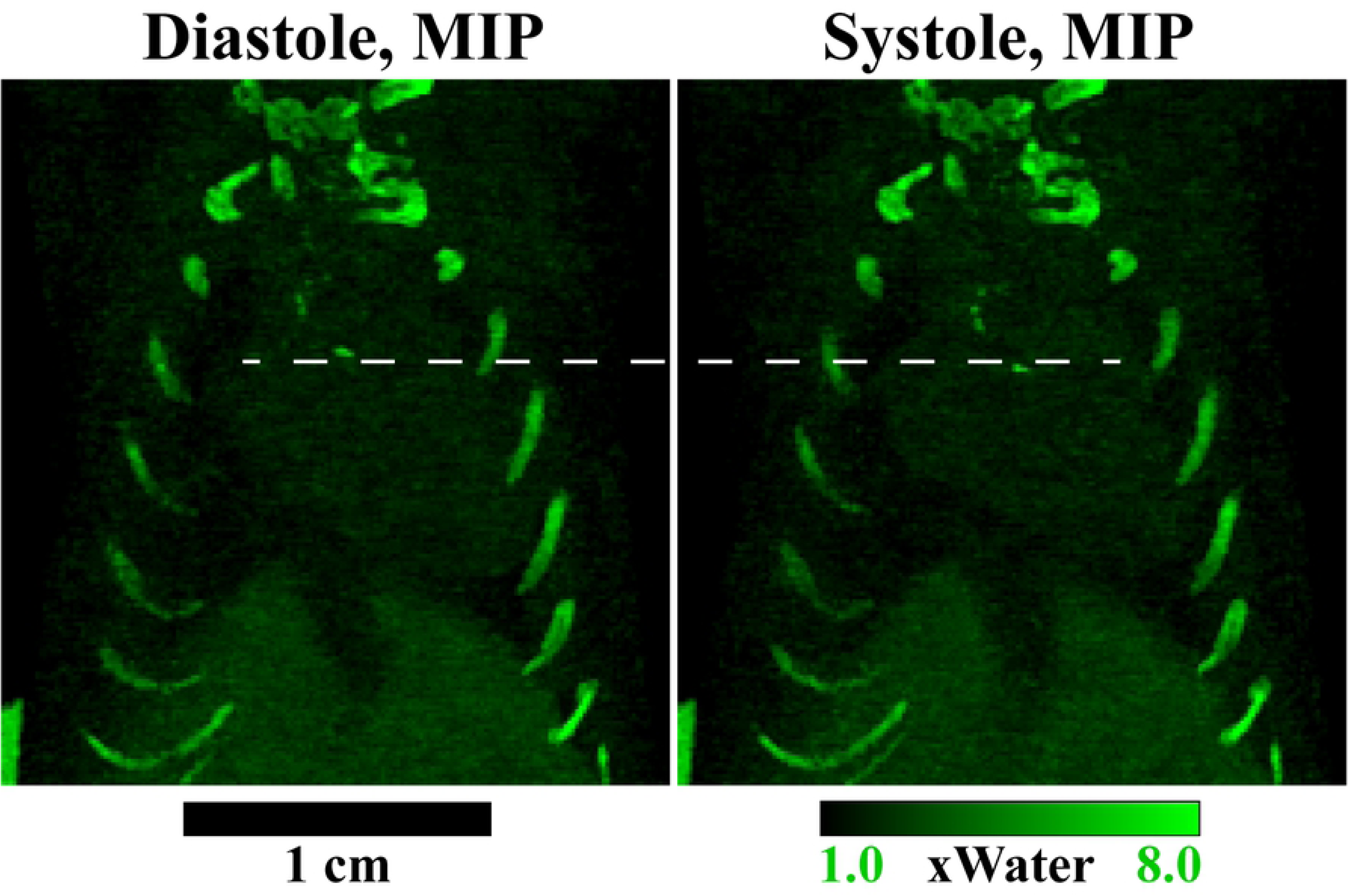
Iterative reconstruction of 4D, PCD-CT data. (A) Algebraic initialization results for each of four energy thresholds: 25-80 keV, 34-80 keV, 40-80 keV, and 55-80 keV. (B) Iterative reconstruction results for the same 2D, coronal slice. (C) Absolute residuals: |(A)-(B)|. Intensity scaling calibration bars (in cm-1) are presented below (A) for (A) and (B), and below (C) for (C). Corresponding noise standard deviation values (measured in water, HU) are shown in the upper left-hand corner of each panel of (A) and (B). Animated, iterative reconstruction results for all cardiac phases are available as supplemental material (S1 Movies).

**Fig 6.**
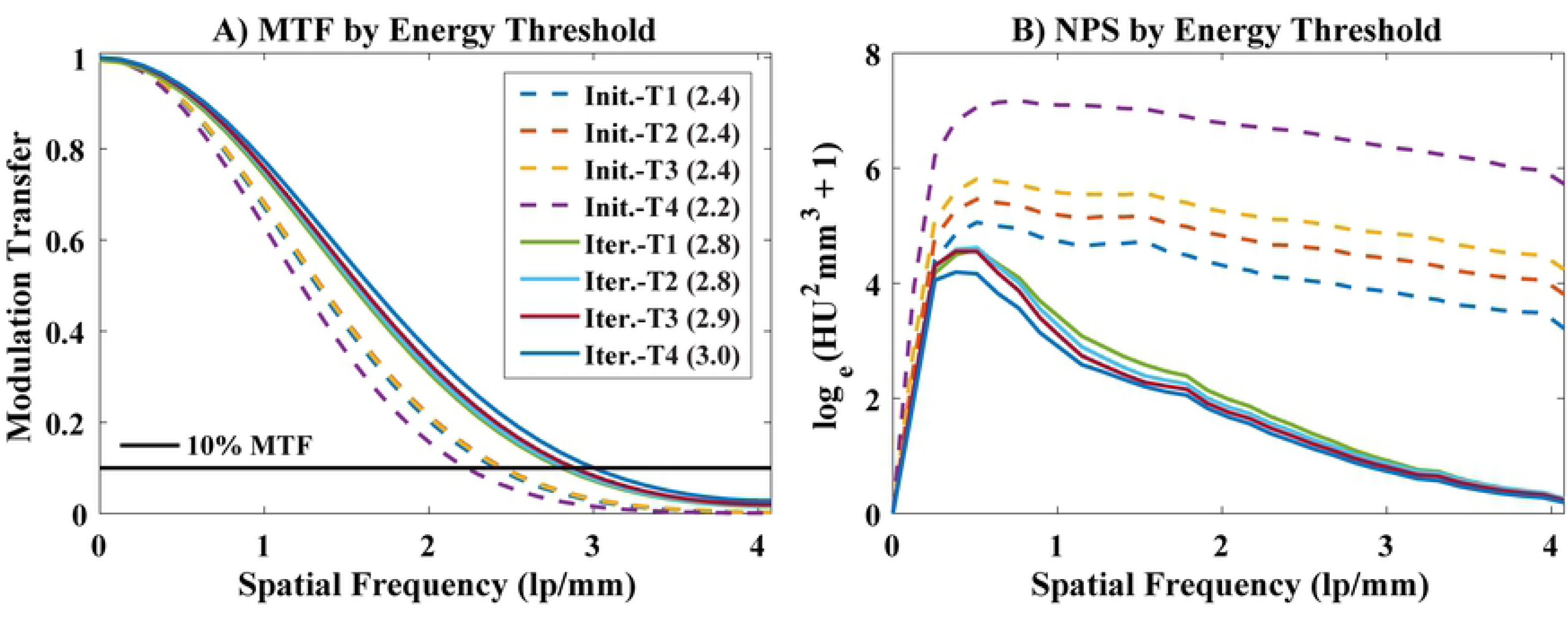
Material decomposition of iterative reconstruction results. Matching 2D slices are shown at ventricular diastole and systole in axial and coronal orientations. Complementary 3D renderings show the four chambers of the heart and the aortic arch, including a segmentation of the left ventricle (LV) used to derive cardiac functional metrics. Decomposition was performed into three basis materials: iodine (red), photoelectric effect (PE, green), and Compton scattering (gray). Calibration vials containing gold (Au, 5 mg/mL), water (W), and iodine (I, 12 mg/mL) can be seen in the axial slices. Calcified plaques appear prominently within the PE map, near the aortic valve and within the aortic arch, and are denoted by dashed circles. Animated material decomposition results are available as supplemental material (S2 Movies).

Fig 6 illustrates the results of image-domain material decomposition applied to the iteratively reconstructed results. The calibration vials in the axial slice indicate effective separation of gold (Au, 5 mg/mL; PE map, green), water (CS map, gray), and iodine (I, 12 mg/mL; red). Furthermore, effective separation is seen between calcium in the bones (PE map) and iodine in the vasculature of the lungs and heart. Noticeably absent are accumulations of gold within the myocardium expected at the site of perfusion defects resulting from radiation-induced myocardial injury. ApoE-/-mice may be less susceptible to radiation-induced myocardial damage than the Tie2Cre;p53^FL/−^ mice, in which endothelial cells are sensitized to radiation, that we studied previously [24]. Yellow, dashed circles in the axial slices and black, dashed circles in the 3D renderings denote the locations of two primary calcifications, one near the aortic valve and one within the aortic arch. Notably, a threshold of 2.5x water in the PE map was used to segment calcifications for the 3D renderings. The 3D renderings also indicate segmentations of the left ventricle (shaded in red).

Fig 7 reiterates the results in this first mouse (Mouse 1, Figs 5-6, 8), and compares them with analogous results in three other mice (10-14 slice maximum intensity projections, MIPs). Calcifications in all mice are denoted by dashed yellow circles, with most of the calcifications seen near the aortic valve and within the aortic arch. Fig 7 also shows the segmented LV volumes by cardiac phase and includes a table of heart and breathing rates, SVs, EFs, and COs measured for each mouse. Overall, our multi-channel, iterative reconstruction is seen to perform robustly across variable heart rates (375-489 beats/min.), breathing rates (124-167 breaths/min.), and ejection fractions (0.42-0.70). Furthermore, comparing attenuation measurements taken in the material vials (Fig 6) included in each of these four mouse scans, iterative reconstruction is seen to robustly preserve spectral contrast relative to the initial algebraic reconstructions. The average spectral bias introduced by iterative reconstruction was measured to be −5.0 HU (over all vials, mice, and energy thresholds). The maximum spectral bias, by magnitude, was 12.2 HU, measured for the fourth energy threshold in the water vial of the Mouse 3 scan.

**Fig 7.**
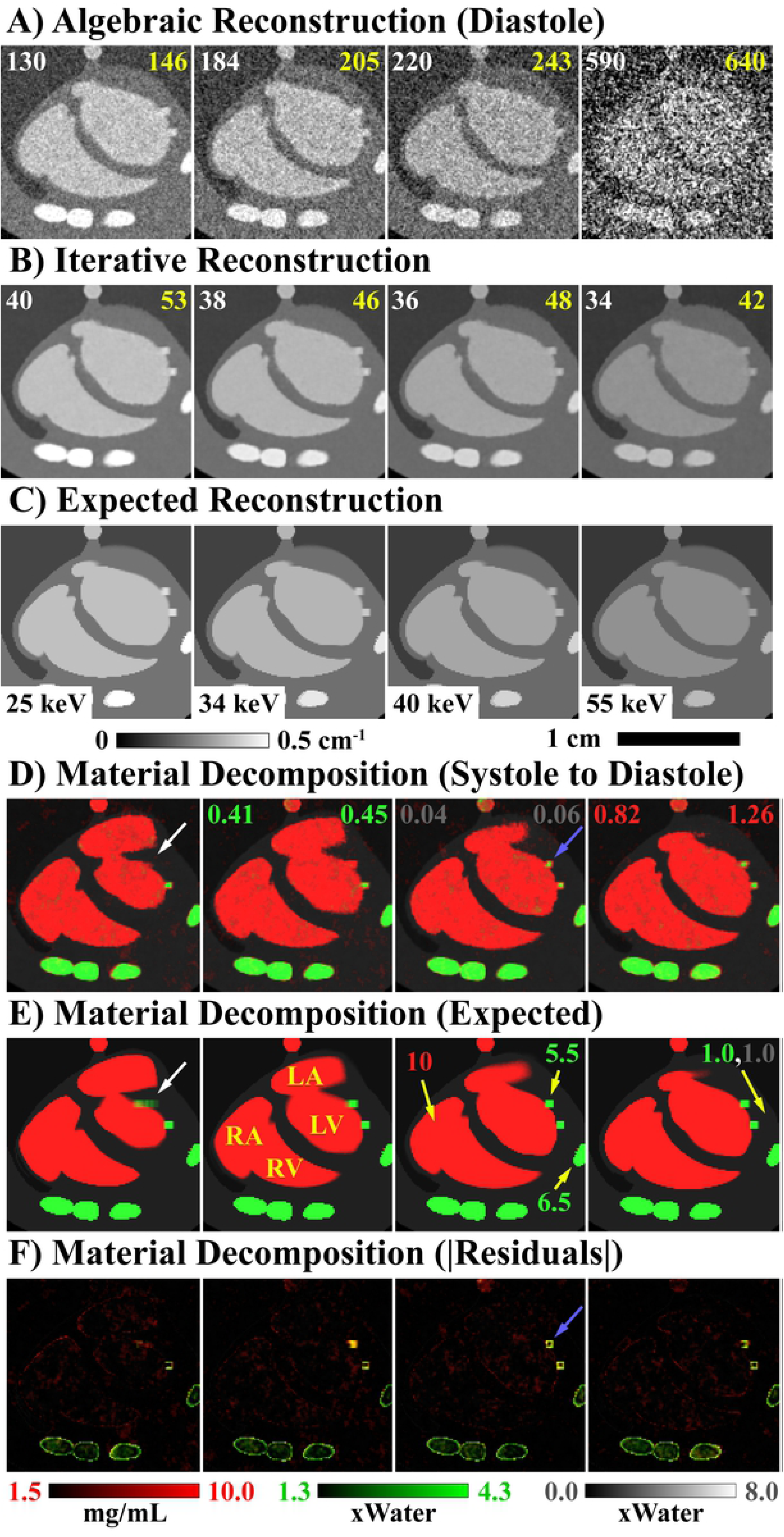
Summary of material decomposition results for data acquired in four mice. Mouse 1, M1, is also shown in Figs 5-6, 8, and the ECG signal for this mouse served as the basis for the MOBY phantom simulation experiment. MIPs are shown through composite material maps (10-14, 2D slices) at ventricular systole (top row) and diastole (bottom row). Calcified plaques are indicated by dashed circles. Animated MIPs are available as supplemental material (S3 Movies). Left ventricular volume curves are plotted for each mouse along with a table of the heart rate (HR, beats/minute), breathing rate (BR, breaths/minute), stroke volume (SV, μL), ejection fraction (EF), and cardiac output (CO, mL/minute) for each mouse.

Finally, Fig 8 focuses on the calcifications indicated in Mouse 1 (Figs 6-7) to illustrate the combined value of gated cardiac imaging and PCD-CT. MIPs through 20 slices of the PE map at diastole and systole highlight the calcifications. By drawing a white, dashed line through identical z planes of diastole and systole, the extent of the motion of the plaque near the aortic valve can be better visualized. Notably, the movement of this calcification over the cardiac cycle is on the order of the size of the calcification (0-4 voxels, 0-500 μm). Combined with the simulation results (Fig 2), this emphasizes the importance of spatial and temporal resolution in accurately localizing and characterizing calcified plaques. More generally, it emphasizes spatial resolution benefits associated with PCD-CT over (dual-source) EID-CT: direct conversion of x-rays to electrons (vs. scintillation) [13] and perfect registration between spectral samples (vs. dual-source geometric calibration).

**Fig 8.**
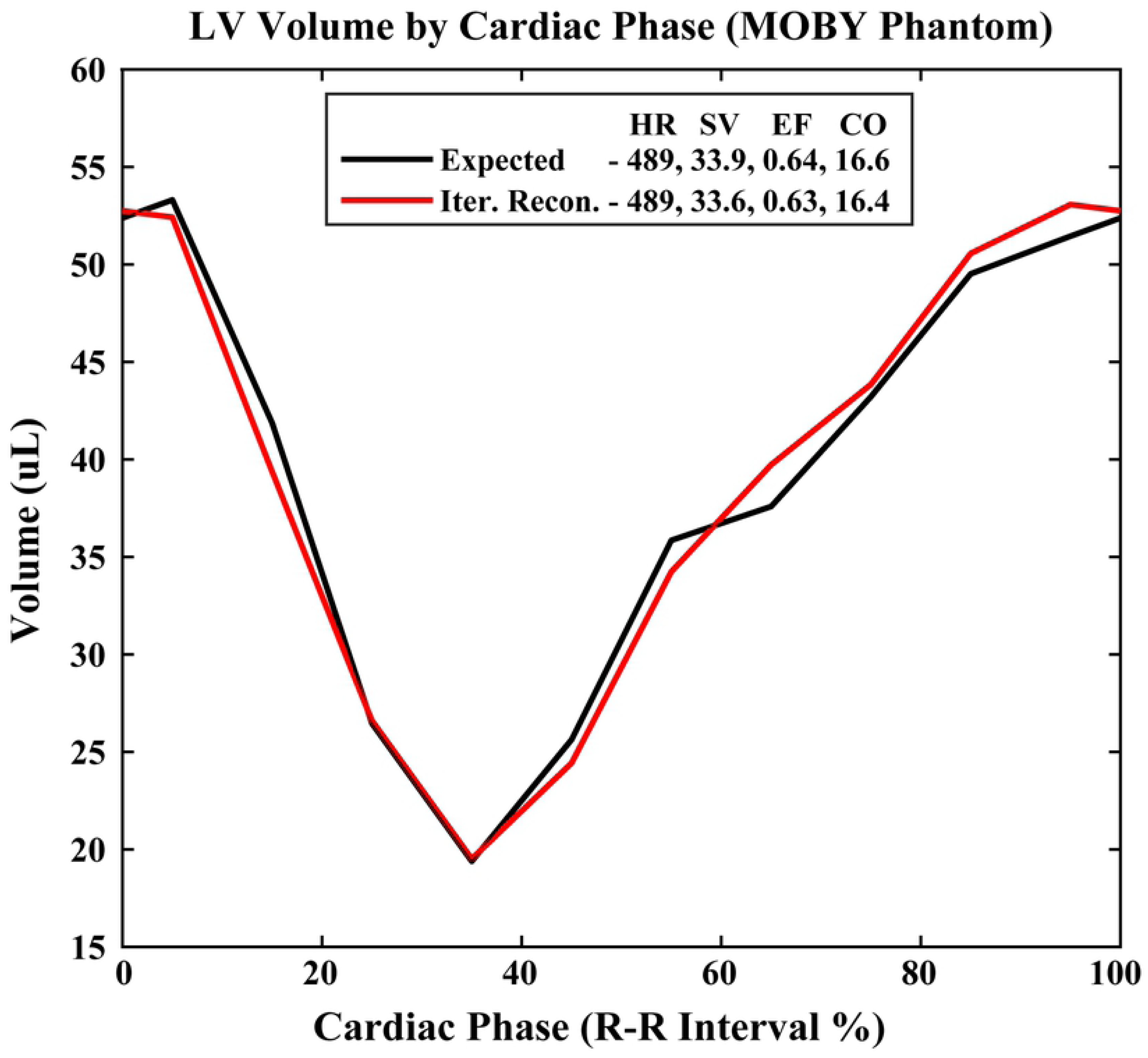
Calcified plaque dynamics. 20-slice MIPs of the PE map at ventricular diastole and systole illustrate the locations of calcifications. A dashed white line indicates an equivalent z position at both phases. Notably, the range of motion of the calcification near the aortic valve is on the order of the plaque dimensions (0-4 voxels, 0-500 μm).

## Discussion and conclusions

In this work, we have demonstrated the feasibility of enhancing *in vivo*, cardiac micro-CT imaging in mice with spectral information provided by a PCD. Given the increased prevalence of high-performance computing hardware, we believe that robust iterative reconstruction methods, such as the one we have demonstrated here, will facilitate the transition from EID-based preclinical micro-CT to PCD-based micro-CT, without associated increases in imaging dose. As illustrated with Fig 8 and in prototype clinical hardware [32], PCDs could provide additional fringe benefits with respect to spatial resolution and the intrinsic registration of spectral data.

Notably, our current methods suffer from common image reconstruction problems. As illustrated in the water phantom (Fig 4), our neighborhood-based regularizers provide limited denoising at low and intermediate spatial frequencies. Furthermore, as illustrated in the MOBY phantom simulation (Fig 2), the ill-conditioning of the material decomposition problem and temporal blurring result in underestimation of calcified plaque sizes in material decomposed images. In extreme cases, calcified plaques may be completely obscured by temporal blurring (Fig 2, white arrows), recommending future work to better characterize implicit tradeoffs between temporal resolution and spectral contrast in the 4D spectral reconstruction problem. We believe that transitioning from neighborhood-based regularizers, like bilateral filtration and RSKR, to multi-scale, convolutional regularizers will be key to the wide-spread adoption of PCD-CT technology and 4D spectral CT. We have already demonstrated that our multi-channel reconstruction framework and convolutional regularizers can reduce 4D, micro-CT reconstruction time by an order of magnitude [33], while also reducing dose and preserving image quality. We are actively working to translate these advancements to the 4D spectral reconstruction problem.

To conclude this work, we acknowledge a growing body of literature which addresses practical issues and limitations associated with current generation PCDs [12, 34–36]. Overcoming sources of spectral distortion such as charge sharing, pulse pileup, K escape, etc., and incorporating more accurate models of PCD data acquisition into the reconstruction process will be equally critical for the long-term adoption of PCD hardware. We look forward to tackling these issues with our peers in future work.

## Acknowledgments

This work was made possible by the loan of a prototype SANTIS 0804 ME photon-counting x-ray detector from DECTRIS, Ltd. (Baden-Dättwil, Switzerland; https://www.dectris.com/). Special thanks to Drs. Spyridon Gkoumas and Thomas Thuering for the installation of the photon counting x-ray detector prototype and for technical support. We also thank Dr. Yi Qi for help with animal support.

## Supporting information

**S1 Movies. Iterative reconstruction results for all cardiac phases.** Video files correspond with the panels of Fig 5B.

**S2 Movies. 2D material decomposition results.** Video files correspond with the axial and coronal 2D images in Fig 6.

**S3 Movies. Material decomposition MIPs from four mice.** Video files correspond with the MIPs in Fig 7.

## References

1. Johnson TR, Krauss B, Sedlmair M, Grasruck M, Bruder H, Morhard D, et al. Material differentiation by dual energy CT: initial experience. European radiology. 2007;17(6):1510–7.

2. Graser A, Johnson TR, Hecht EM, Becker CR, Leidecker C, Staehler M, et al. Dual-energy CT in patients suspected of having renal masses: can virtual nonenhanced images replace true nonenhanced images? Radiology. 2009;252(2):433–40.

3. Chae EJ, Song J-W, Seo JB, Krauss B, Jang YM, Song K-S. Clinical utility of dual-energy CT in the evaluation of solitary pulmonary nodules: initial experience. Radiology. 2008;249(2):671–81.

4. Yu L, Leng S, McCollough CH. Dual-energy CT–based monochromatic imaging. American journal of Roentgenology. 2012;199(5_supplement):S9–S15.

5. Johnson T, Fink C, Schönberg SO, Reiser MF. Dual energy CT in clinical practice: Springer Science & Business Media; 2011.

6. Clark DP, Ghaghada K, Moding EJ, Kirsch DG, Badea CT. In vivo characterization of tumor vasculature using iodine and gold nanoparticles and dual energy micro-CT. Physics in Medicine and Biology. 2013;58(6):1683.

7. Ashton JR, Clark DP, Moding EJ, Ghaghada K, Kirsch DG, West JL, et al. Dual-energy micro-CT functional imaging of primary lung cancer in mice using gold and iodine nanoparticle contrast agents: a validation study. PLOS ONE. 2014;9(2):e88129.

8. Mukundan S Jr, Ghaghada KB, Badea CT, Kao C-Y, Hedlund LW, Provenzale JM, et al. A liposomal nanoscale contrast agent for preclinical CT in mice. American Journal of Roentgenology. 2006;186(2):300–7.

9. Clark DP, Badea CT. Hybrid spectral CT reconstruction. PLOS ONE. 2017;12(7):e0180324.

10. Cruje C, Holdsworth DW, Gillies ER, Drangova M. High-concentration gadolinium nanoparticles for pre-clinical vascular imaging. Medical Imaging 2018: Physics of Medical Imaging: International Society for Optics and Photonics; 2018. p. 105732N.

11. Shikhaliev PM. Energy-resolved computed tomography: first experimental results. Physics in Medicine & Biology. 2008;53(20):5595.

12. Taguchi K, Iwanczyk JS. Vision 20/20: Single photon counting x-ray detectors in medical imaging. Medical Physics. 2013;40(10):100901.

13. Willemink MJ, Persson M, Pourmorteza A, Pelc NJ, Fleischmann D. Photoncounting CT: technical principles and clinical prospects. Radiology. 2018;289(2):293–312.

14. Symons R, Krauss B, Sahbaee P, Cork TE, Lakshmanan MN, Bluemke DA, et al. Photon-counting CT for simultaneous imaging of multiple contrast agents in the abdomen: an in vivo study. Medical physics. 2017;44(10):5120–7.

15. Cormode DP, Si-Mohamed S, Bar-Ness D, Sigovan M, Naha PC, Balegamire J, et al. Multicolor spectral photon-counting computed tomography: in vivo dual contrast imaging with a high count rate scanner. Scientific reports. 2017;7(1):4784.

16. Gao H, Yu H, Osher S, Wang G. Multi-energy CT based on a prior rank, intensity and sparsity model (PRISM). Inverse Problems. 2011;27(11):115012.

17. Candès EJ, Li X, Ma Y, Wright J. Robust principal component analysis? Journal of the ACM (JACM). 2011;58(3):11.

18. Clark DP, Lee C-L, Kirsch DG, Badea CT. Spectrotemporal CT data acquisition and reconstruction at low dose. Medical Physics. 2015;42(11):6317–36.

19. Holbrook M, Clark D, Badea C. Low-dose 4D cardiac imaging in small animals using dual source micro-CT. Physics in Medicine & Biology. 2018;63(2):025009.

20. Meganck JA, Liu B. Dosimetry in micro-computed tomography: a review of the measurement methods, impacts, and characterization of the Quantum GX imaging system. Molecular Imaging and Biology. 2017;19(4):499–511.

21. Badea CT, Clark DP, Holbrook M, Srivastava M, Mowery Y, Ghaghada KB. Functional imaging of tumor vasculature using iodine and gadolinium-based nanoparticle contrast agents: a comparison of spectral micro-CT using energy integrating and photon counting detectors. Physics in medicine and biology. 2019.

22. Piedrahita JA, Zhang SH, Hagaman JR, Oliver PM, Maeda N. Generation of mice carrying a mutant apolipoprotein E gene inactivated by gene targeting in embryonic stem cells. Proceedings of the National Academy of Sciences. 1992;89(10):4471–5.

23. Zhang SH, Reddick RL, Piedrahita JA, Maeda N. Spontaneous hypercholesterolemia and arterial lesions in mice lacking apolipoprotein E. Science. 1992;258(5081):468–71.

24. Lee C-L, Min H, Befera N, Clark D, Qi Y, Das S, et al. Assessing cardiac injury in mice with dual energy-microCT, 4D-microCT, and microSPECT imaging after partial heart irradiation. International Journal of Radiation Oncology* Biology* Physics. 2014;88(3):686–93.

25. Clark D, Badea C. Data-efficient methods for multi-channel x-ray CT reconstruction. Medical Imaging 2018: Physics of Medical Imaging: International Society for Optics and Photonics; 2018. p. 105732A.

26. Clark DP, Badea CT. Joint regularization for spectro-temporal CT reconstruction. Proceedings of SPIE Medical Imaging 2016. p. 1–11.

27. Clark DP, Badea CT, editors. GPU-Based Tools for Multi-Channel X-ray CT Reconstruction. The Fifth International Conference on Image Formation in X-Ray Computed Tomography; 2018; Salt Lake City, Utah.

28. Alvarez RE, Macovski A. Energy-selective reconstructions in x-ray computerised tomography. Physics in Medicine and Biology. 1976;21(5):733.

29. Segars W, Tsui B. 4D MOBY and NCAT phantoms for medical imaging simulation of mice and men. Journal of Nuclear Medicine. 2007;48(supplement 2):203P–P.

30. Yushkevich PA, Piven J, Hazlett HC, Smith RG, Ho S, Gee JC, et al. User-guided 3D active contour segmentation of anatomical structures: significantly improved efficiency and reliability. Neuroimage. 2006;31(3):1116–28.

31. Siewerdsen JH, Wojciech Z, Xu J. Chapter 4: Cone-beam CT image quality. In: Shaw CC, editor. Cone beam computed tomography: Taylor & Francis; 2014. p. 37–58.

32. Leng S, Rajendran K, Gong H, Zhou W, Halaweish AF, Henning A, et al. 150-μm Spatial Resolution Using Photon-Counting Detector Computed Tomography Technology: Technical Performance and First Patient Images. Investigative radiology. 2018;53(11):655–62.

33. Clark D, Badea C. Convolutional regularization methods for 4D, x-ray CT reconstruction. Medical Imaging 2019: Physics of Medical Imaging: International Society for Optics and Photonics; 2019. p. 109482A.

34. Taguchi K, Stierstorfer K, Polster C, Lee O, Kappler S. Spatio-energetic cross-talk in photon counting detectors: Numerical detector model (Pc TK) and workflow for CT image quality assessment. Medical physics. 2018;45(5):1985–98.

35. Yu Z, Leng S, Li Z, Halaweish AF, Kappler S, Ritman EL, et al. How low can we go in radiation dose for the data-completion scan on a research whole-body photon-counting CT system. Journal of computer assisted tomography. 2016;40(4):663.

36. Ballabriga R, Campbell M, Heijne E, Llopart X, Tlustos L, editors. The Medipix3 prototype, a pixel readout chip working in single photon counting mode with improved spectrometric performance. 2006 IEEE Nuclear Science Symposium Conference Record; 2006: IEEE.

